# Predicting the responses in colour matching experiments from the symmetries of colour space

**DOI:** 10.1101/2023.09.16.557954

**Authors:** Nicolás Vattuone, Inés Samengo

## Abstract

In conceptual spaces, the distance between concepts is represented by a metric that cannot usually be expressed as a function of a few, salient physical properties of the represented items. For example, the space of colours can be endowed with a metric capturing the degree to which two chromatic stimuli are perceived as different. As many optical illusions have shown, the colour with which a stimulus is perceived depends, among other contextual factors, on the chromaticity of its surround, an effect called “chromatic induction”. Heuristically, the surround pushes the colour of the stimulus away from its own chromaticity, increasing the salience of the boundary. Previous studies have described how the magnitude of the push depends on the chromaticity of both the stimulus and the surround, concluding that the space of colours contains anisotropies and inhomogeneities. The importance of contextuality has cast doubt on the practical or predictive utility of perceptual metrics, beyond a mathematical curiosity. Here we provide evidence that the metric structure of the space of colours is indeed useful and has predictive power. By using a notion of distance between colours emerging from a subjective metric, we show that the anisotropies and inhomogeneities reported in previous studies can be eliminated. The resulting symmetry allows us to derive a universal curve for the average chromatic induction that contains no fitting parameters and is confirmed by experimental data. The theory also predicts the magnitude of chromatic induction for every possible combination of stimulus and surround demonstrating that, at least in the case of colours, the metric captures the symmetries of perception, and augments the predictive power of theories.

## 1 Introduction

In trichromats, all perceivable colours can be specified by three coordinates. The collection of all triplets of numbers that represent the entire gamut of perceivable colours conforms a set, but not necessarily a space. In order for a set to be upgraded to a fully-fledged space additional structure is required. In a Riemmanian space, for example, the additional structure is provided by a metric. Riemman himself postulated that there might be a suitable metric for colour space [25]. Since then, many metrics have been proposed, some of grounded in theoretical principles [27, 24, 44, 5, 6], and others from experimental data [39, 40, 19, 32, 38], among others.

Recently, the Riemannian nature of colour space was questioned [3]. The objection was based on the principle of diminishing returns, which states that the perceived difference between two distant colours is less than the sum of the distances of each of them to an intermediate colour lying along the geodesic connecting them. However, the experimental evidence supporting this claim was collected along a curve in colour space that can be argued not to be a geodesic. First, the CIELab coordinates are not perceptually uniform [26, 18]. Second, the employed stimuli were surrounded by a different chromaticity, which affects the perceived colour even when stimulus and surround have markedly different chromaticities [38].

In line with this sceptical view of the Riemmanian nature of the space of colours, Fairchild [10] questioned the utility of conceiving the set of colours as a *space*, by casting doubt on whether the additional structure required to turn a set into a space reflects a property of perception. Taking Fairchild’s example, even though diamonds are usually described by four numbers, “clarity”, “colour”, “cut” and “weight”, the quadruplets formed from these coordinates by no means define a space. Admittedly, in the realm of conceptual mental representations, rather daring choices of coordinates have sometimes been postulated as potentially useful spaces, such as the ones spanned by the length of the legs and necks of birds [4], or the weight of a car and the power of its engine [1]. Therefore, assessing what exactly is gained, beyond an aesthetic satisfaction, by conceiving the set of colours as a Riemannian space is a legitimate question.

The present paper provides evidence for the claim that, at least in certain domains, it is indeed useful to endow a conceptual space with a metric. We worked with the space of colours as an example, and showed that the perceptual metric that stems from just noticeable differences embodies the symmetries of the internal representation of colours. Consequently, chromatic induction, in the perceptual coordinates, can be described as a homogeneous and isotropic process. We believe that it is this symmetry, above all, what justifies the promotion of the set of colours to a space of colours. Moreover, if the perceptual coordinates are known, the symmetry hypothesis makes it possible to predict the results of certain experiments on chromatic induction without fitting parameters and for a wide range of experimental setups. Therefore, even chromatic induction, which was so far believed to vary in strength from point to point in colour space, becomes amenable to a simple theoretical description.

*Chromatic induction* is the phenomenon by which the colour of a stimulus is altered by the chromaticity surrounding it [12]. For example, a stimulus perceived as grey in a spatially uniform setting acquires a bluish tint when surrounded by yellow. Surrounds have long been known to exert a repulsive effect [36, 42, 37, 41, 9], because the alteration they produce in the colour of the stimulus occurs along a direction that enhances the perceptual difference between stimulus and surround. In the example above, yellow is more different from blue than grey. Importantly, the direction of the repulsive shift is not exactly complementary [17], and its magnitude has been shown to depend in non-trivial ways on the chromaticities of both the stimulus and the surround [15]. Such complex dependencies have led the scientific community to believe that chromatic induction is non homogeneous and non isotropic throughout colour space. Here we show that the asymmetries disappear when endowing the space of colours with an appropriate metric, and that the previously collected experimental data can be reconciled with a symmetric model of chromatic induction that is uniform throughout colour space.

In a previous paper, we showed the existence of a set of colour coordinates, in which the perceptual displacement exerted by a given surround on each possible stimulus can be represented as a repulsive field centred at the surround that is both isotropic and homogeneous (Fig. 1). The property of isotropy is evident in the field’s rotational symmetry, which means that (a) the surround shifts the colour of the stimulus along a straight line originating at the surround and passing through the stimulus, and (b) the magnitude of the displacement solely depends on the Euclidean distance between the stimulus and the surround. The property of homogeneity implies that when the surround changes, the repulsive field shifts rigidly to the new location, with no distortions. In this paper we show that the symmetric theory can be reconciled with the asymmetric experiments with a coordinate transformation between these special coordinates to the more widely employed experimentally. The transformation maps the straight lines in Fig. 1 into curved lines in the experimental coordinates, making the observed anisotropies compatible with a symmetric induction. The underlying symmetry can be used to predict a hidden regularity in the experimental results. In other words, although in the collected data the magnitude of chromatic induction was reported to be different for each combination of stimulus and surround, the proposed metric explains those differences away, as resulting from the specific choice of coordinates.

**Figure 1.**
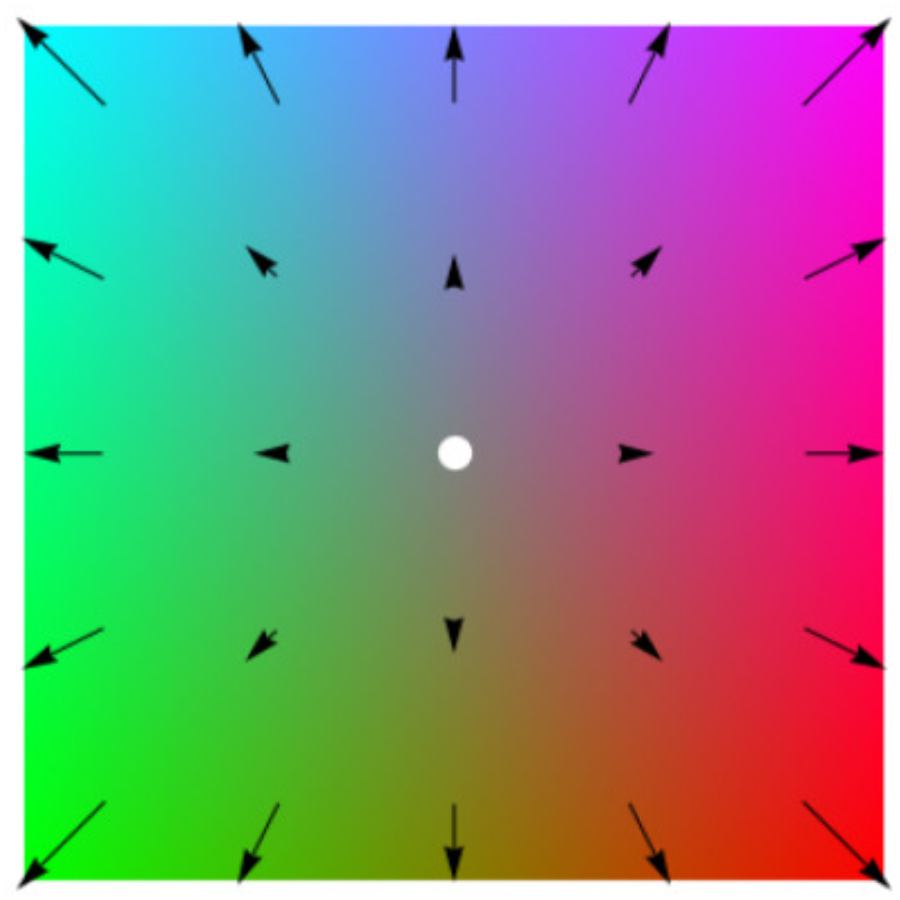
Chromatic induction as a displacement in colour space. The location of the white dot indicates the chromaticity of the surround (grey, in this example). The repulsive field illustrates how this surround modifies the colour with which different stimuli are perceived. The base of each arrow indicates the chromaticity of a stimulus, and the tip marks the chromaticity of another stimulus that, when shown in a spatially uniform setting, is perceived as equal to the stimulus at the base of the arrow, when the latter is enclosed by the chosen surround. We hypothesise that in the perceptual coordinates, the field is isotropic, that is, it has rotational symmetry. We also hypothesise that the space is homogeneous, by assuming that, when the chromaticity of the surround varies, the field shifts rigidly to a new location.

## 2 Methods

In Sect. 2.1, the space of colours is defined as the collection of classes of equivalence that can be obtained from spatially structured matching experiments. In Sect. 2.2, we define a notion of distance in colour space that reflects the subjective notion of similarity. This metric is determined by the discrimination thresholds obtained from experiments in which an observer compares two very similar chromaticities. The so called *perceptual coordinates*, as well as the transformations that leave the metric invariant are introduced in Sect. 2.3. The Methods section closes with a brief description of the experiments used to characterise the perceptual effects, the symmetry of which is assessed in this paper.

### 2.1 Colors as classes of equivalence

A *colour* is a private experience that emerges from the perception of a luminic stimulus. Experiments have established that there is no one-to-one mapping of the colour of a stimulus with its spectral properties. Here we use the term *chromaticity* to indicate the 3-dimensional description of the spectral properties of the stimulus to which most trichromat humans are sensitive. Chromaticity is typically reported in coordinate systems such as *SML, RGB, XY Z*, etc. The colour of a stimulus cannot be paired with its chromaticity because the context in which the stimulus is displayed plays an important role in the perceptual experience, as illustrated by the example of chromatic induction. Let *E* define a set of experimental conditions that include both the chromaticity *x* of a stimulus and the chromaticity *b* of its surround: *E* = (*x, b*). Even though many properties of the experimental setting are known to exert an effect on how the stimulus is perceived [14, 21, 30], for simplicity, in this paper we work with spatially uniform surrounds, and fixate all other contextual variables, so that *x* and *b* suffice to specify the experimental conditions. In this case, 6 coordinates determine the pair *E* = (*x, b*): 3 for *x* and 3 for *b*. Luckily, a remarkable simplifying property of visual perception comes into play, namely, that for every stimulus *x*^1^ surrounded by chromaticity *b*^1^, and for a given chromaticity *b*^2^, a stimulus of chromaticity *x*^2^ exists, such that the colour of stimulus *x*^2^ surrounded by *b*^2^ is perceived as equal to the colour *x*^1^ surrounded by *b*^1^. To formally state this property, we introduce the notation *x* ⫽ *b* to represent a stimulus of chromaticity *x* enclosed by a uniform surround of chromaticity *b*. The perceptual equality of *x*^1^ ⫽ *b*^1^ with *x*^2^ ⫽ *b*^2^ is represented as *x*^1^ ⫽ *b*^1^ ∼ *x*^2^ ⫽ *b*^2^. The simplifying porperty stated above can now be restated as: For every stimulus *x*^1^ surrounded by chromaticity *b*^1^, and for a given chromaticity *b*^2^, a stimulus of chromaticity ***x***^2^ exists, such that *x*^1^ ⫽ *b*^1^ ∼ *x*^2^ ⫽ *b*^2^. This property motivated Resnikoff [24] to define the space of colours as the set of equivalence classes generated by the matching operation among stimuli with different surrounds (see also Berthier [2] and Provenzi [23]). The colour instantiated by *x*⫽ *b* and all those in the same class of equivalence is denoted [*x* ⫽ *b*]. This quotienting operation defines a 3-dimensional space of colours.

We assume that, at least for the unsaturated chromaticities employed in experiments performed with LED monitors, all equivalence classes contain a unique *uniform representative*, that is, a pair of the form *x* ⫽ *x* with its surround. We define the function **Φ**_*b*_(*x*) as the one that maps each member *x* ⫽ *b*, in which the stimulus coincides *b* of a given class to its uniform representative *x*^0^ ⫽ *x*^0^, such that

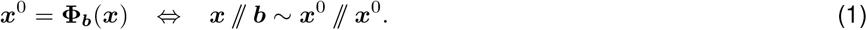

We call **Φ**_*b*_(*x*) the *induction function* since it encodes the information of how the perception of the stimulus with chromaticity *x* is altered when changing from the setup *x* ⫽ *x* to *x* ⫽ *b*.

### 2.2 Perceptual distance

Many of the interesting properties of physical space are rooted in geometric relations. For instance, the isotropy and homogeneity of physical laws stems from the invariance of Euclidean distance under rotations and translations. In Vattuone et al. [38], we showed that also in colour space a perceptual geometry exists, in which the symmetries of perception are revealed. The geometry can be constructed empirically from the results of discrimination experiments in which the just noticeable differences (JNDs) of colour space are determined. Of course, such results may vary from observer to observer. The derived perceptual distance between two colours is a measure of how dissimilar they are perceived.

Since colours are classes of equivalence in the space of pairs *x* ⫽ *b*, a distance between colours can only depend on the class, and not on the individual values of *x* and *b*. Following Vattuone et al. [38], to define the distance explicitly, we use the notion of uniform representative: The distance *d*(*α, β*) between two classes of equivalence *α* and *β* can be reported as a function of the chromaticities *x*^*α*^ and *x*^*β*^ of the corresponding uniform representatives, that is,

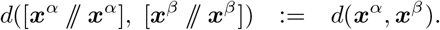

The procedure to explicitly obtain this distance from experimentally measured JNDs is developed in [38].

### 2.3 Perceptual coordinates and symmetries in the metric

In several reported discrimination experiments [16, 38], the space of colours was revealed to be flat, or compatible with a flat geometry [19, 5, 6]. In other words, the metric obtained from discrimination experiments was shown to have nearly zero curvature. In flat spaces, a coordinate system exists in which the perceptual distance coincides with the Euclidean distance. We call such coordinates *perceptual*. The transformation that maps the chromaticities of uniform representatives (expressed, for example, as cone fundamentals *SML*) onto the perceptual coordinates can be parametrised by a few numbers that may vary from observer to observer.

In the perceptual coordinates the metric is Euclidean, so the transformations that leave the metric invariant are rotations and translations. Since the metric was derived from the JNDs obtained from discrimination experiments, by construction, in the perceptual coordinates all JNDs remain constant throughout colour space, and along all the directions in which the difference is to be detected. The question remains, however, whether other perceptual properties also remain invariant under rotations and translations. Here we explore this topic in the case of chromatic induction. Induction is isotropic and homogeneous if [38]

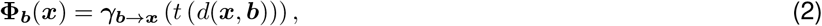

where *γ*_*b*→*x*_ is the geodesic in colour space joining [*b* ⫽ *b*] and [*x* ⫽ *x*]. In the perceptual coordinates, this geodesic is a straight line, so the shift induced by the surround is radially oriented. In Eq. 2, *d* is the Euclidean distance, and *t* is a function that depends only on the distance between the stimulus and the surround and is equal to *t*(*d*(*x, b*)) := *d*([*b* ⫽ *b*], [*x* ⫽ *b*]).

### 2.4 Setup and coordinates in the colour matching experiment

A scheme of the experimental protocol of Klauke and Wachtler [15] is shown in Fig. 2A. A bipartite field was displayed on the screen, both halves with the same luminance. One of the halves, the so-called *achromatic*, was coloured with the reference grey *g*: 40 *cd/m*^2^, *CIE*[*x, y*] = [0.310, 0.316]. The chromaticity *c* of the other half was fixed to different values throughout the experiment. The left-right disposition of the two halves was switched randomly from trial to trial. Inside the field of chromaticity *c*, a 2° wide square patch with chromaticity *r* (the *reference* stimulus) was shown. Inside the achromatic field, a patch of equal size was displayed, the chromaticity *s* of which was adjusted by the subject to maximise the similarity between the two patches.

**Figure 2.**
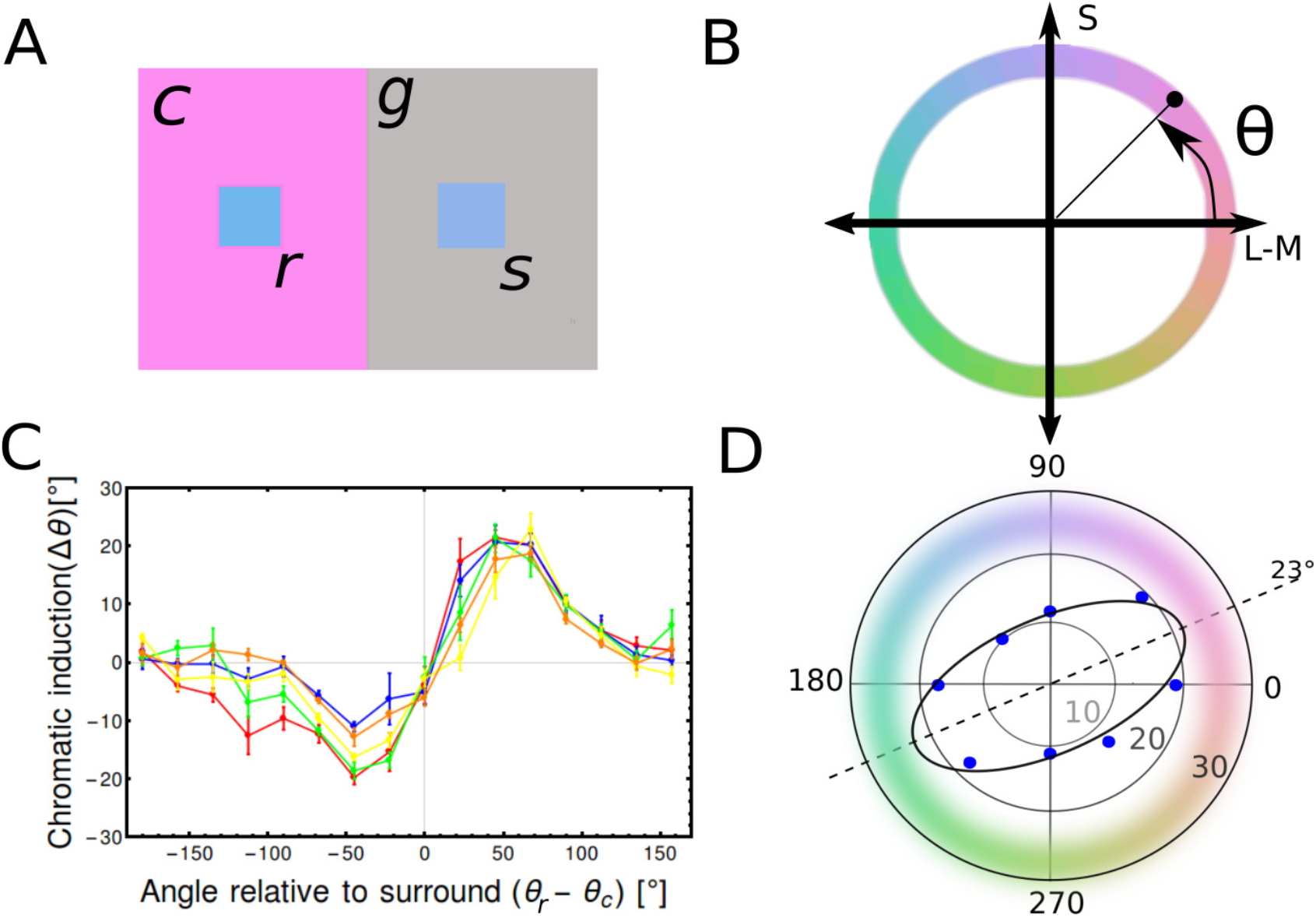
Experimental setup and main results of Klauke and Wachtler [15]. **A**: On one side of the screen a reference square of chromaticity ***r*** was displayed enclosed by a uniform surround of chromaticity ***c***. On the other side, a second square was surrounded by an achromatic stimulus *g*. The observer adjusted the chromaticity ***s*** of the patch surrounded by grey so as to maximise its similarity with the patch surrounded by ***c*. B**: The search for the matching chromaticity was performed along a chromatic circumference defined on the iso-luminant plane generated by the *S* and *L* − *M* coordinates of *DKL* space [8]. The angle *θ* was defined with respect to the *L* −*M* axis. **C**: Chromatic induction (defined as the angular difference between the selected ***s*** and the reference *r* chromaticities) obtained for different observers and plotted as a function of the angular difference *θ*_*r*_ −*θ*_***c***_ between the reference stimulus and the chromatic surround, for a pink surround (*θ*_***c***_ = 0°). **D**: Polar plot of the anisotropy of chromatic induction. The distance to the origin is the maximal chromatic induction (expressed in degrees), as measured for each of the 8 surrounds *c*, corresponding to the angles 0°, 45°, …, 315°. The major axes of the ellipse points along the surround for which induction is maximal. Data points were taken from [15].

The cone-contrast coordinates are functions of the *Cone Fundamentals* [33] 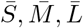

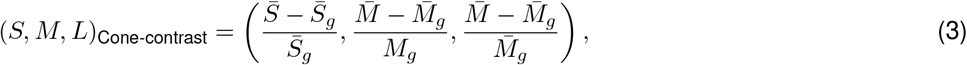

where the sub-index *g* denotes the reference grey. Klauke and Wachtler’s experiment was performed on an isoluminant plane that for each subject was determined through heterochromatic flicker photometry [34, 13]. The results were reported in opponent cone-contrast *DKL* coordinates [8], in which the *S*-axis represent chromaticities of varying 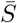 value, while simultaneously adjusting 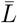 and 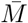 to keep the luminance constant. The orthogonal axis contained in the iso-luminant plane is named *L* − *M*, and represents the difference of the corresponding cone contrast coordinates.

On the isoluminant plane, stimuli were identified by their chroma, or saturation 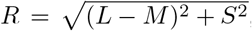, and their hue *θ* = ArcTan(*S/*(*L* − *M*)). The locus along which the matching stimulus was searched for was defined by the equation

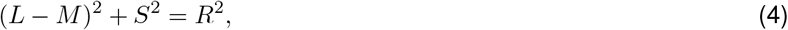

with a fixed value for *R* (Fig. 2 B). The locus was parametrised as *L* − *M* = *R* cos(*θ*) and *S* = *R* sin(*θ*). The experiment tested 8 different chromatic surrounds *c*, each combined with 16 reference stimuli *r*, both uniformly distributed in the angular coordinate *θ*. All the tested surrounds *c* had the same luminance as the neutral gray *g*, and their saturation was 20% lower than that of the locus containing the tested and responded stimuli *r* and *s*.

## 3 Results

### 3.1 Experimental characterization of chromatic induction

The psychophysical tasks used to characterise induction [15] consist of asymmetric matching experiments, in which each subject is requested to adjust the chromaticity *s* of a patch enclosed by a grey surround of chromaticity *g* so as to maximise the perceptual similarity with a second patch of given reference chromaticity *r*, which was in turn enclosed by a surround of chromaticity *c* (Fig. 2). In order to avoid an exceedingly lengthy protocol, the search is typically done along a locus of fixed chroma *R* and varying hue *θ* (Sect. 2.4). In the *DKL* isoluminant plane, chroma and hue correspond to polar coordinates, and the locus is shaped as a circumference (Fig. 3A). Klauke and Wachtler [15] showed that induction followed a fairly universal sine-like behaviour (Fig. 2C), similar to the tilt illusion with oriented bars [11]. Confirming previous studies [12, 29, 41], the sign of the induction, defined as Sign(*θ*_*s*_ − *θ*_*r*_), coincided with the sign of the angular difference *θ*_*r*_ − *θ*_*c*_, supporting the repulsive nature of induction. Starting from the spatially uniform condition *r* = *c*, as the reference stimulus *r* began to deviate from its surround *c*, the magnitude of the shift *θ*_*s*_ − *θ*_*r*_ increased, until it reached a maximum at a certain reference stimulus *r*_m_, and then decreased gradually, to vanish at *θ*_*r*_ = *θ*_*c*_ + 180°. The angular position *r*_m_ of the maximum always differed from *θ*_*c*_ in less than 90°, and maximal induction never exceeded 20°.

**Figure 3.**
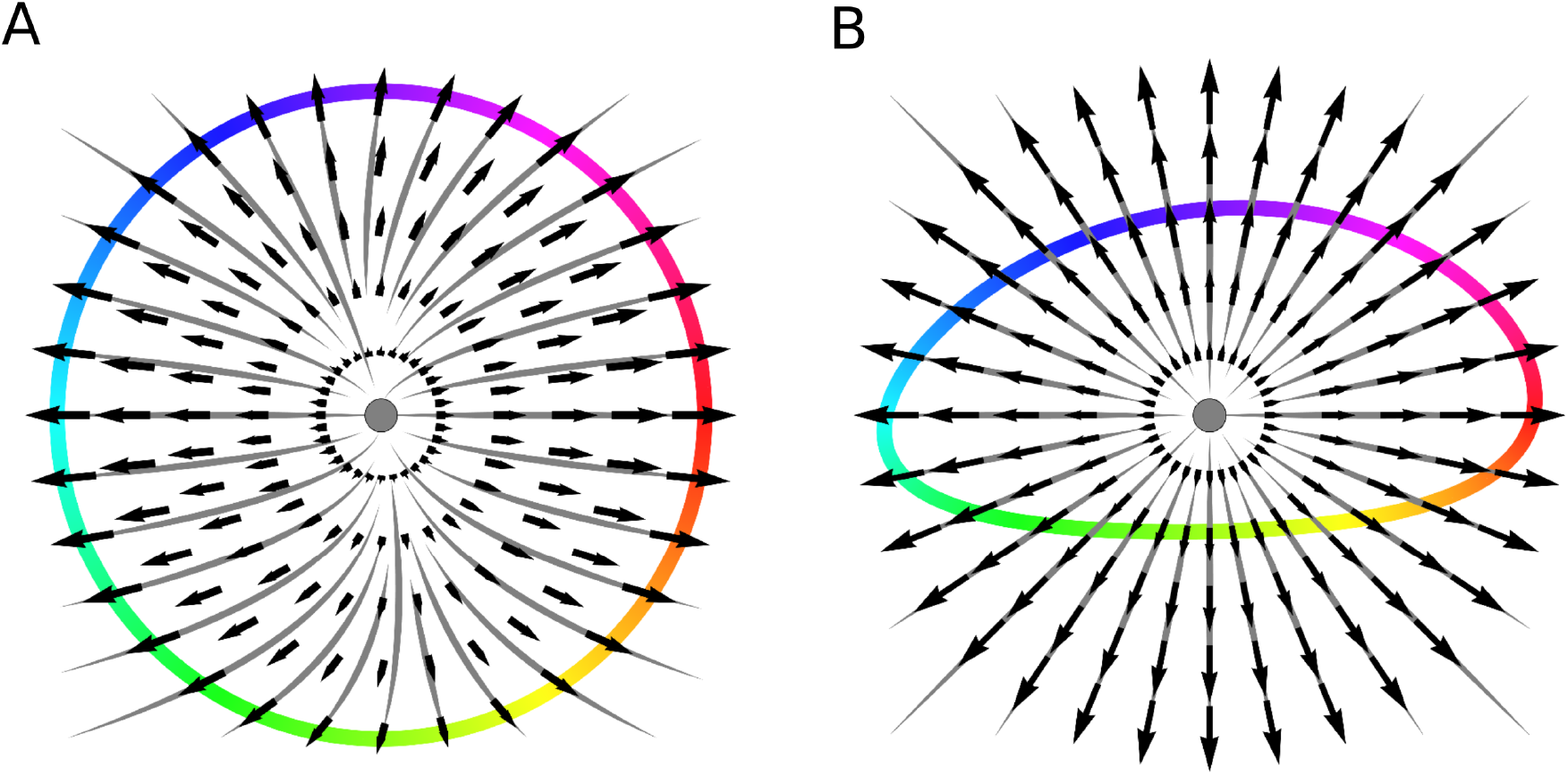
Experimental setup of the asymmetric matching experiment [15] experiment. (A) In *DKL* coordinates, the locus containing the candidate matching stimuli has rotational symmetry, but the field representing induction does not. (B) In perceptual coordinates, the field representing induction has rotational symmetry, but the locus appears as flattened.

Klauke and Wachtler [15] also showed that the shape of the curve changed when the colour *c* of the surround was varied (Sect. 3.4.1). These changes, though small, were systematic across different subjects. Therefore, although qualitatively speaking all surrounds produced similar sine-like induction curves, a significant anisotropy could be detected at the quantitative level. In Fig. 2 D, the maximal induction *θ*_*s*_ − *θ*_*r*_ obtained at *r*_m_ is displayed in a polar plot as a function of the angular position *θ*_*c*_ of the surround. Induction was maximal at *θ*_*c*_ ≈ ±23°, corresponding to magenta or greenish surrounds, and bluish or yellowish reference patches. A differential mechanism in the processing of colours associated with daylight statistics was hypothesised to underlie the anisotropy, as a putative evolutionary advantage of subjects adapted to their natural environment [43, 7]. Here we explore the extent up to which a fully symmetric description of induction can still account for this observed anisotropy.

The locus in which the matching stimulus was searched for is defined by Eq. 4. A circumference is the set of points that lie all at the same distance from a given central point. Therefore, Eq. 4 only defines a circumference if the pair (*L* − *M, S*) denotes two Cartesian coordinates in an Euclidean space. However, if the notion of distance between colours is intended to represent a perceptual distance, then the *DKL* coordinates are not Euclidean, since previous experiments have shown examples of pairs of colours that all lie at the same *DKL* Euclidean distance, and yet, are discriminated with unequal difficulty (see the different sizes and excentricities of the discrimination ellipses reported by Krauskopf and Gegenfurtner [16], and confirmed by Vattuone et al. [38]). In a non-Euclidean space, the locus of Eq. 4 is not a circumference (Fig. 3B). A spherically symmetric induction field may then well produce a perceptual shift whose dependence on the difference *θ*_*r*_ − *θ*_*c*_ varies with the surround *θ*_*c*_, since along the tested locus, the perceptual shift has no rotational symmetry. Here we analyse the hypothesis that the observed anisotropy stem from the mismatch between the symmetry of the locus employed in the study and the symmetry of the perceptual shift produced by induction (Fig. 3).

### 3.2 Analytical description of chromatic induction

#### 3.2.1 General expression

In this section, we analytically derive the magnitude and direction of the inductive shift produced by chromatic surrounds in experiments where the shape of the locus is arbitrary. We are interested in working with curves of unconstrained shape to account for the fact that a change of coordinates from the original *DKL* space, where the locus is circular, to a new set, where induction is symmetric, may change the curve substantially. We assume that a set of coordinates exists, the so-called *perceptual* coordinates (Sect. 2.3), in which the perceptual distance between two colours *x* = (*x*_1_, *x*_2_) and *y* = (*y*_1_, *y*_2_) is captured by the Euclidean formula

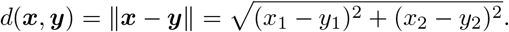

The fundamental hypothesis is that in these coordinates, induction can be described by a spherically symmetric repulsive field, as postulated in Eq. 2 and illustrated in Fig. 1. The curve **𝒞** (*θ*) in which the matching stimulus is searched for is only assumed to be smooth and periodic, that is, **𝒞** (0) = **𝒞** (2*π*).

In the perceptual coordinates, the locus can be parametrised as **𝒞** (*θ*) = (**𝒞** _1_(*θ*), **𝒞** _2_(*θ*)). The reference stimulus *r* belongs to the locus, so an angle *θ*_*r*_ exists, such that *r* = **𝒞** (*θ*_*r*_). When enclosed by the surround *c*, the reference chromaticity produces the colour [**𝒞** (*θ*_*r*_) *c*] (Sect. 2.1). On the other hemi-field, the stimuli *s* = **𝒞** (*θ*_*s*_) available to the subject also belong to the locus, and since they are enclosed by a grey surround *g*, each one of them produces the percept [**𝒞** (*θ*_*s*_) *g*]. The subject adjusts the value of *θ*_*s*_ to maximise the perceptual similarity between the two colours, or equivalently, to minimise the distance

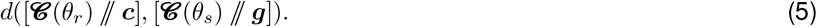

The minimisation condition is weaker than the perceptual equality *r* ⫽*c* ∼ *s* ⫽*g*. The latter can only be fulfilled if the matching stimulus ***s*** belongs to the locus, in which case, the minimal distance vanishes. Since the locus is defined by the experimental setup, the equality is by no means guaranteed to be attainable [31]. Whenever a perfect match is not possible, the subject is instructed to select the colour that best approximates the match, that is, the one minimising the perceptual distance. This distance can be written as

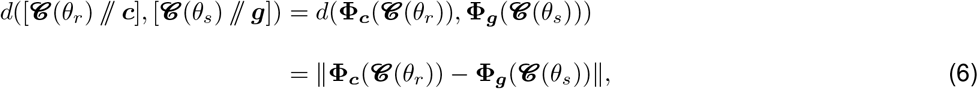

where **Φ**_*c*_(*x*) is the spatially uniform chromaticity that is perceptually equal to *x* ⫽ *c* (Sect. 2.1). Calculating the derivative with respect to *θ*_*s*_ and equating it to zero, at the selected *θ*_*s*_,

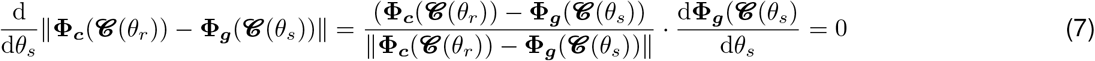

The functions **Φ**_*c*_ and **Φ**_*g*_ represent the percepts induced by the backgrounds *c* and *g*, respectively. Equation 7 implies that the minimal distance is achieved when the difference **Φ**_*c*_(**𝒞** (*θ*_*r*_))− **Φ**_**g**_(**𝒞** (*θ*_*s*_)) is orthogonal to the locus, since the vector d**Φ**_***g***_(**𝒞** (*θ*_*s*_))*/*d*θ*_*s*_ is tangent to the curve at the colour *s* ⫽ *g*. A realisation of this minimisation is represented in (Fig. 4A).

**Figure 4.**
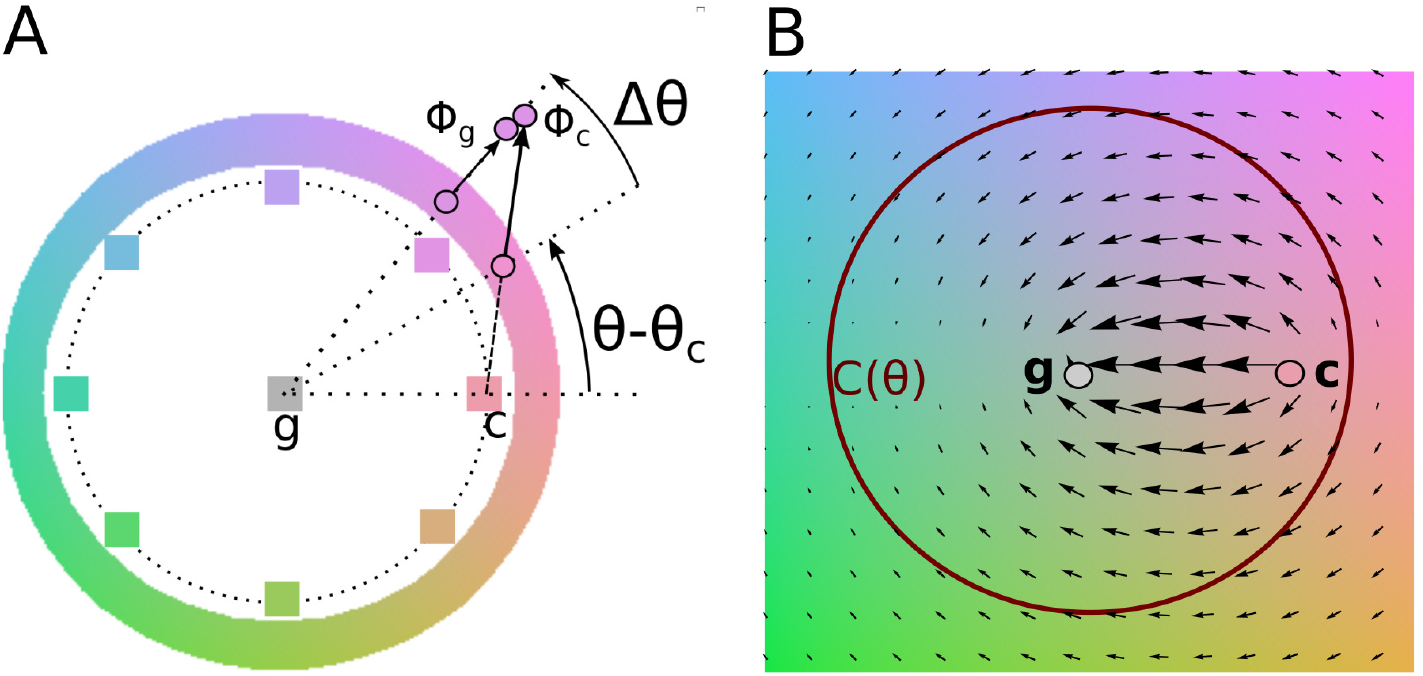
(A) On one of the hemi fields, the grey surround *g* displaces the adjustable stimulus *s* along the direction *s* − *g*. In turn, on the other hemi field, the surround *c* displaces the reference stimulus *r* along the direction *r* − *c*. Both shifts are here greatly enlarged, for visualisation purposes. The perceptual match is produced when the displaced percepts **Φ**_*g*_(*s*) and **Φ**_***c***_(*r*) coincide. (B) When all the points on the curve **𝒞** (*θ*_*r*_) are sufficiently far away from the grey surround *g*, the stimulus *s* that best approximates the match is the one for which the projection of the dipolar field on the curve has the value predicted by Eq. 11.

Experiments show that perceptual displacements Δ*θ* = *θ*_*s*_ − *θ*_*r*_ are small, with an absolute maximum no larger than 20°*≈* 0.35 rad. At first order in Δ*θ*, Eq. 7 becomes

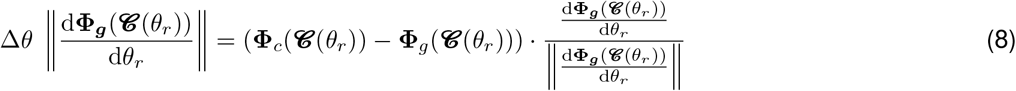

If the locus **𝒞** (*θ*) and the function **Φ**_***b***_(***x***) are known, this equation provides a simple way to calculate Δ*θ*. In the following section, we show that under suitable assumptions for **Φ**_***b***_(***x***), the displacement Δ*θ* depends only on **𝒞** (*θ*), with only a single scaling parameter characterising the only relevant property of the inductive field **Φ**_***b***_(***x***): its asymptotic strength.

#### 3.2.2 Small induction and fast saturation approximation

Experiments have shown that the deviation of **Φ**_*b*_(*x*) from *x* grows monotonically as *x* departs from the surround *b* [38]. The growth, however, saturates fairly rapidly, so in this section, we assume that the perceptual shift undergone by stimuli that are moderately different from their surrounds are governed by the asymptotic behaviour of **Φ**_***b***_(*x*). Under this premise, Eq. 8 can be simplifed, so that Δ*θ* becomes independent of other properties of **Φ**_***b***_(*x*). The derivation used here is valid whenever the locus (*θ*) always remains sufficiently far from the two surrounds *g* and *c*, so that the inductive displacements **Φ**_***g***_(*s*) and **Φ**_***c***_(*r*) can be assumed to have reached the maximal magnitude, a condition that is not difficult to attain by experimental setups in which the locus is more saturated than the surrounds. As a result, Eq. 8 is transformed to an expression that, except for a single numerical constant that can be obtained experimentally, no longer depends on the functional shape of **Φ**_***a***_(*b*). The perceptual displacement Δ*θ*, hence, becomes a function of the shape of the locus **𝒞** (*θ*) alone.

If the surround has no influence on the perceived colour, then **Φ**_***b***_(*x*) = *x*. If there is an influence, as a first approximation it is reasonable to assume that the intensity of chromatic induction has a maximum value 𝓀, and experiments confirm that this value is smaller than 20^deg^ [15, 38]. We decompose

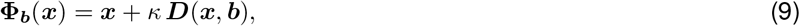

where 𝓀*D*(*x, b*) is the perceptual displacement of the stimulus *x* produced by the surround *b*, and ⫽*D*⫽ ≤ 1. The hypothesis that **Φ**_***b***_(*x*) can be assumed to be symmetric (Sect. 2.3) implies that in perceptual coordinates, *D* is oriented radially and has rotation invariance,

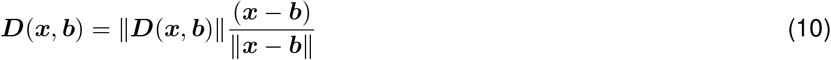

In the perceptual coordinates, the assumption that **Φ**_***b***_(*x*) saturates rapidly implies that a characteristic distance 𝓁 exists for which ⫽D(*x, b*) ⫽ ≈ 1 whenever ⫽*x* − *b*⫽ is larger than 𝓁.

We set the origin of the coordinate system at the reference gray: *g* ≡ (0, 0). If all the points of the locus **𝒞** (*θ*) lie at a distance larger than 𝓁 from the surround *c*, Eq. 8 becomes

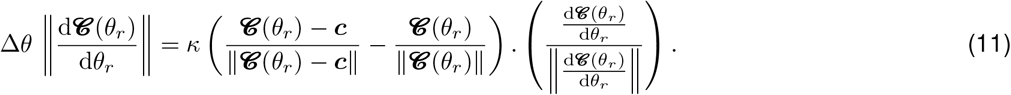

This equation specifies the value of the angular displacement Δ*θ* for an arbitrary chromatic curve **𝒞** (*θ*). Except for the scale factor *κ*, which is the only parameter that describes the intensity of induction and must be fitted experimentally, the result only depends on the geometry of the curve.

Equation 11 has an easy interpretation. On the left hand side, 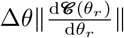 represents the differential arc length of the displacement. The right hand side contains the difference of two radial fields, one of them centred at the neutral grey *g* = 0 and the other at the chromatic surround *c*. Both have unit strength, so their difference produces a dipolar field as in Fig. 4B. The field is projected on the vector tangent to the curve **𝒞** (*θ*), because the perpendicular component does not change the choice of the subject, due to the restriction imposed by the locus. In what follows, Eq. 11 is referred to as the *approximate model*.

### 3.3 Chromatic induction for circular curves

In this section, we use Eq. 11 to calculate the inductive shift obtained for a locus **𝒞** (*θ*) that is circular. We know that the curves used in experiments so far are not circular — at least not in the perceptual coordinates. Yet, the circular case is a good starting point for a later expansion in increasing degrees of deviation from circularity.

We first calculate Δ*θ* using the approximate model in the case in which **𝒞** (*θ*) is a circumference of radius *R* parametrised by **𝒞** (*θ*) = *R* (cos *θ*, sin *θ*). Expressing the surround *c* in polar coordinates as *c* = *c* (cos *θ*_*c*_, sin *θ*_*c*_) and replacing this expression in Eq. 11, we get

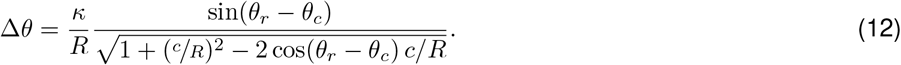

The induction described by Eq. 12 is isotropic, since the displacement Δ*θ* only depends on the angular difference *θ*_*r*_ − *θ*_*c*_, which remains invariant under global rotations. The only free parameter is the ratio *κ/R*, which acts as a scale factor determining the maximal strength of induction. The reference colour ***r*** for which induction is maximal is located at the angle 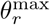 that fulfills the condition

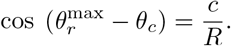

Maximal induction is obtained for a reference colour that differs from the surround in no more than *π/*2. When the locus passes far away from the surround 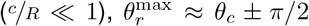. Instead, if the locus passes right through the surround (*R* = *c*), maximal induction is obtained for reference colours that coincide with their surround, that is, 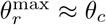. This case, however, is not compatible with the assumption ⫽*x* − *b*⫽ ≫ 𝓁 used to derive Eqs. 11 and 12.

The ratio 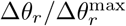 between the displacement for the reference colour *r* and the displacement for the reference colour for which induction is maximal has no fitting parameters. In Fig. 5, this ratio (black line) is compared with the results obtained by Klauke and Wachtler [15], for which *c/R* = 0.8. For this value, Eq. 12 predicts that the maximal induction occurs at *θ*_*r*_ = *θ*_*c*_ ± 36°. The experiments were performed for several surrounds, namely *θ*_*c*_ = 0°, ±45°, ±90°, ±135°, 180°. Although due to the non-circularity of the locus, the analytical result of Eq. 5 does not describe the inductive shift obtained with any individual surround, quite surprisingly, it does capture the behaviour of the data when the latter are averaged across all the 8 surrounds. In Sect. 3.4.4 we derive this result. But before that, in the following section, we develop an analytical description of the curve Δ*θ* corresponding to each surround, considering paths **𝒞** (*θ*) that progressively deviate from the circular case.

**Figure 5.**
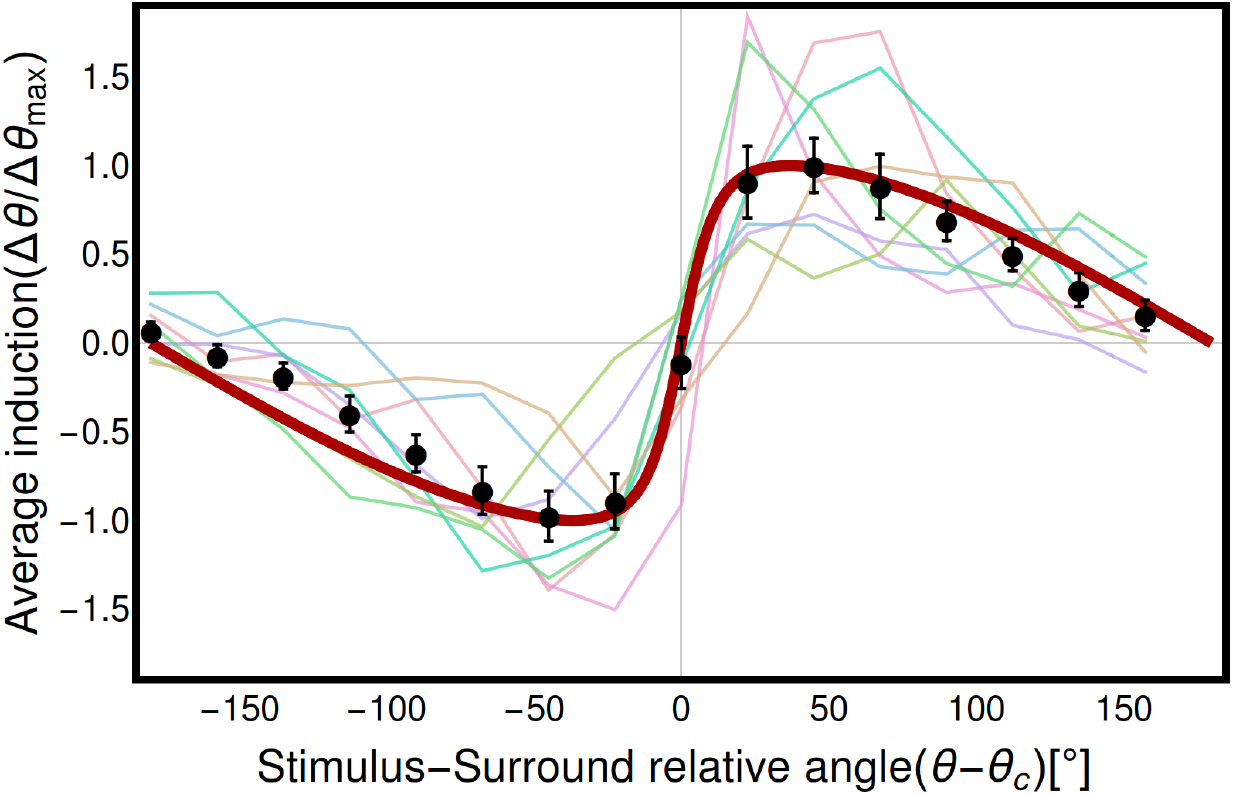
Thick red line: Theoretical scaled angular displacement produced by induction when the locus **𝒞** (*θ*) is circular. Scaling consists of dividing the analytical result of Eq. 12 by the maximal value of induction along the locus. Equation 12 is also the mean displacement obtained for non-circular curves, when averaged among surrounds ***c***, valid when the departure of the circular case is small (Sects. 3.4.4 and 5.1). Thin coloured lines: Experimental displacements obtained for the non-circular locus employed by Klauke and Wachtler [15] for each tested surround. These curves were all scaled by the same constant, such that the maximum of their average was equal to 1. Black dots with error bars: Average and standard error of the mean of the 8 thin coloured lines.

### 3.4 Apparent anisotropy for non-circular loci

In the perceptual coordinates, JNDs remain constant throughout colour space. Asymmetric matching experiments have so far been performed in other coordinate systems, for example *DKL*, in which JNDs vary from place to place, and from tested direction to tested direction. For example, Krauskopf and Gegenfurtner [16] observed larger JNDs when the tested differences were performed along the *S* axis than along the *L* − *M* axis. To obtain more uniform JNDs, they scaled the *S* axis appropriately. As a consequence, the locus **𝒞** (*θ*), which appeared to be circular in the *DKL* coordinates, became elliptic in the scaled ones. In Sect. 3.4.1, we derive the inductive displacement Δ*θ* expected for elliptic loci.

More generally, the required scaling could perhaps involve the stretching or contraction of an axis that does not coincide with any of the cardinal axes of MacLeod and Boynton [20]. In Sect. 3.4.2, we extend the derivation of the inductive displacement Δ*θ* to encompass these cases as well. There we see that the additional freedom provided by the possibility of scaling along an arbitrary direction improves the predictability of the experimentally observed displacements.

A still more comprehensive scenario is one in which the relation between the experimental coordinates and the perceptual ones is nonlinear. Indeed, Krauskopf and Gegenfurtner [16] showed that the JNDs along the *S* axis were not only in general larger than those along the *L* − *M*, but also, that they increased with *S*. A nonlinear transformation to the perceptual coordinates distorts the locus **𝒞** (*θ*) even beyond an ellipse. In Sect. 3.4.3 we derive the inductive displacements Δ*θ* when the locus employed in the experiment is described by a curve that contains up to a 4^th^ power of the coordinates. This additional freedom does not yield an improvement in the explanation of the experimental data. Importantly, all three extensions beyond the circular locus result in inductive displacements Δ*θ* that are no longer a function of *θ*_*r*_ − *θ*_*c*_. In other words, anisotropic results are a hallmark of a transformation between the experimental and the perceptual coordinates that goes beyond a mere scaling.

Finally, in Sect. 3.4.4 we derive the inductive shift for a locus of arbitrary shape. We demonstrate that, as long as the deviation from the circular case is small, when the curve Δ(*θ*) is averaged in all the surrounds *θ*_*c*_, Eq. 12 is recovered (red curve of Fig. 5), implying that the anisotropies cancel out, and the circular result prevails.

#### 3.4.1 Elliptic 𝒞 aligned with the coordinate axes

In several previous studies, discrimination thresholds have been reported to be elliptic, and the ellipses had principal axes aligned with the cardinal directions of the *DKL* space [16, 22]. Therefore, a first approximation of the perceptual coordinates (*x*_1_, *x*_2_) can be constructed by scaling one of the two axes by a factor *β*, that is, 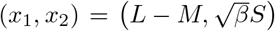 In perceptual coordinates thus constructed, a chromatic locus that appeared to be circular in *DKL* coordinates, becomes an ellipse with eccentricity 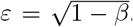. The inductive shift Δ*θ* can be calculated analytically from Eq. 11. If the value of *ε* is know from discrimination experiments, the shape of Δ*θ* can be obtained with no fitted parameters. If the value of *ε* is unknown, it may be fitted from the experimentally obtained shifts. This we have done with the data collected by Klauke and Wachtler [15], and obtained *ε* = 0.84 ± 0.03. With the fitted *ε*, the coloured curves in Fig. 6 are derived. The obtained fit is acceptable (*χ*^2^ = 1.27 per degree of freedom, Table 2), and it represents a significant improvement from the fit of that same experimental data with just the model for a circular locus (*χ*^2^ = 1.94 per degree of freedom). Quite remarkably, fitting a single parameter *β* suffices to predict the way Δ*θ*(*θ*_*r*_) varies with all 8 tested surrounds *θ*_*c*_. In particular, the reference stimulus 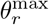 for which induction is maximal is well predicted by the theory (Fig. 7).

**Table 1:**
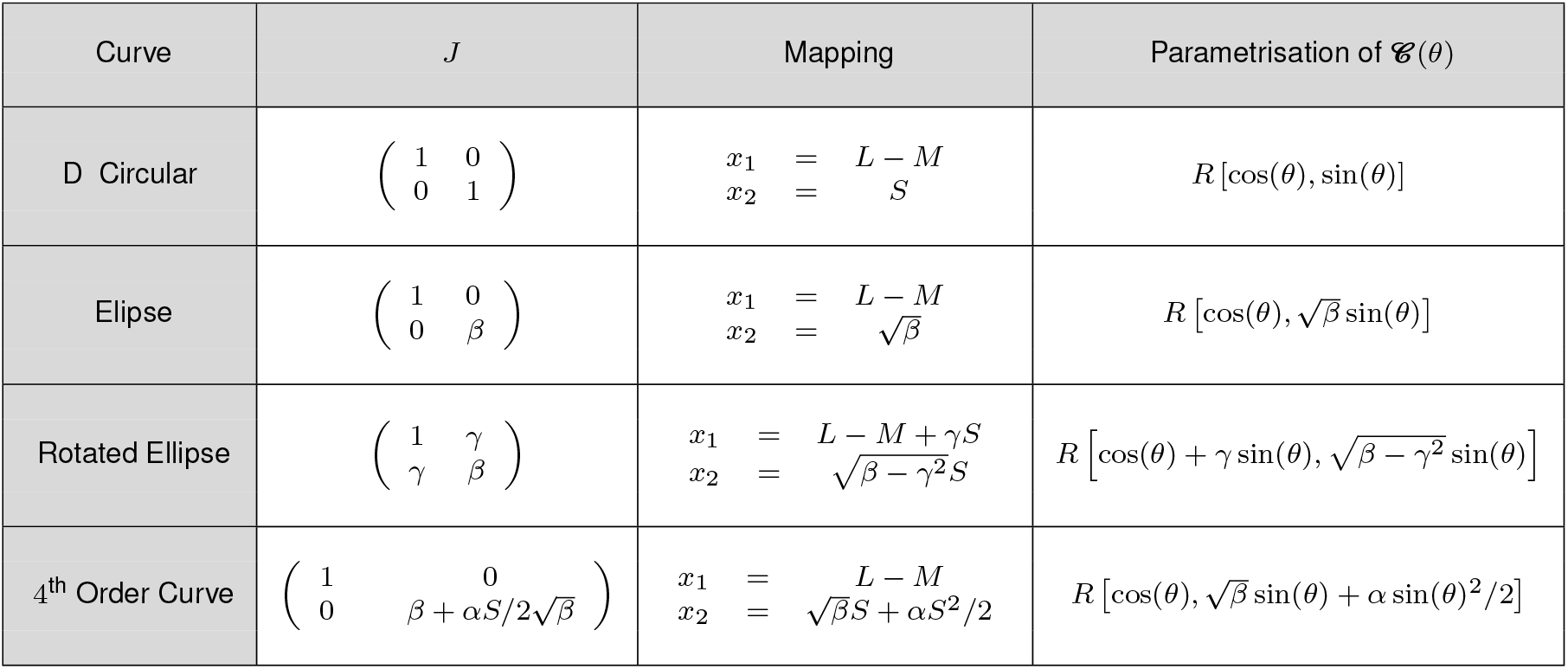
Summary of the studied models, in increasing level of complexity. The matrix *J* is the quadratic form representing the ellipses of the *JND* of detection experiments. The mapping specifies the transformation from the *DKL* coordinates to the perceptual coordinates that turn *J* into the unit matrix. The parametrisation assumes a circular curve in the original *DKL* coordinates, and describes the shape of the curve in the perceptual ones.

**Table 2:** Values of the parameters that, for each model, best fit the data. *χ*^2^*/N*_*df*_ is the ratio of the *χ*^2^ value and the number of degrees of freedom. For the sake of completeness, the parameters *β* and *γ*, which are not free parameters but functions of ϵ and *ϕ*_0_ are also included.

| Model | $\kappa/R(10^{-2})$ | $\ell/R$ | $\varepsilon$ | $\varphi_0(^{\circ})$ | $\alpha$ | $\beta(\epsilon, \varphi_0)$ | $\gamma(\epsilon, \varphi_0)$ | $\chi^2/N_{df}$ |
| --- | --- | --- | --- | --- | --- | --- | --- | --- |
| Circular (Approximate) | $2.2 \pm 0.1$ | - | - | - | - | - | - | 1.94 |
| Circular (Complete) | $2.3 \pm 0.1$ | $0.16 \pm 0.10$ | - | - | - | - | - | 1.94 |
| Ellipse (Approximate) | $1.83 \pm 0.08$ | - | $0.84 \pm 0.03$ | - | - | $0.29 \pm 0.04$ | - | 1.27 |
| Ellipse (Rotated) | $1.73 \pm 0.07$ | - | $0.88 \pm 0.02$ | $-13 \pm 2$ | - | $0.27 \pm 0.03$ | $-0.18 \pm 0.02$ | 0.963 |
| Ellipse (Control) | $2.6 \pm 0.2$ | - | $0.6 \pm 0.2$ | $8 \pm 8$ | - | $0.6 \pm 0.2$ | $0.05 \pm 0.05$ | 1.80 |
| Ellipse (Complete) | $1.75 \pm 0.07$ | $0.08 \pm 0.04$ | $0.88 \pm 0.02$ | $-14 \pm 2$ | - | $0.27 \pm 0.03$ | $-0.18 \pm 0.02$ | 0.953 |
| 4<sup>th</sup> Order (Approximate) | $1.84 \pm 0.08$ | - | $0.83 \pm 0.03$ | - | $0.12 \pm 0.05$ | $0.31 \pm 0.04$ | - | 1.23 |
| 4<sup>th</sup> Order (Rotated, complete) | $1.76 \pm 0.07$ | $0.07 \pm 0.04$ | $0.88 \pm 0.02$ | $-14 \pm 2$ | $0.04 \pm 0.04$ | $0.27 \pm 0.03$ | $-0.18 \pm 0.02$ | 0.955 |

**Figure 6.**
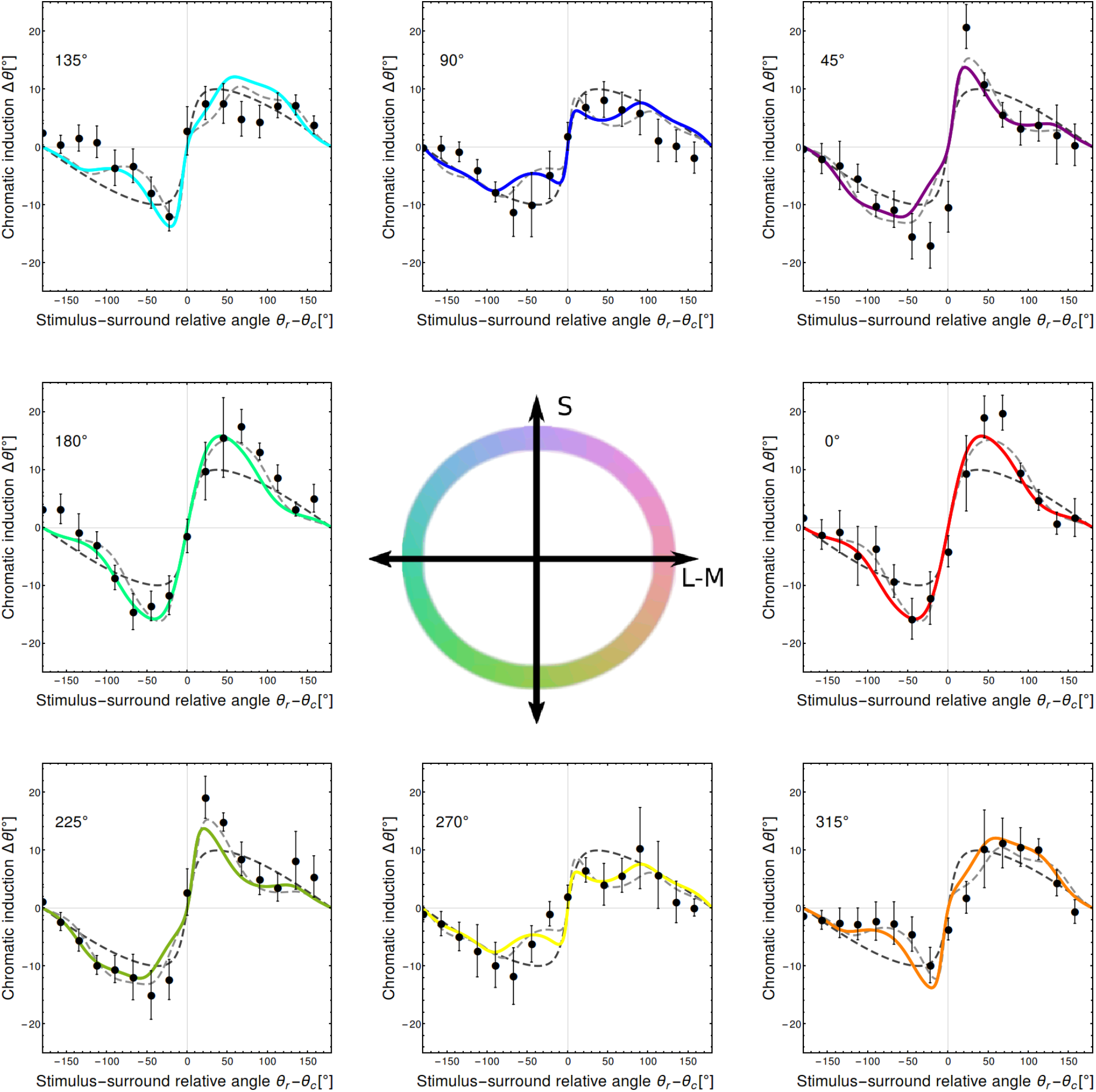
Chromatic induction for eight different surrounds (data points re-drawn from Klauke and Wachtler [15]). Each panel shows the angular displacement Δ*θ* = *θ*_*s*_ − *θ*_*r*_ between the responded and the reference stimuli as a function of the angle *θ*_*r*_ − *θ*_*c*_. Points indicate population mean (*N* = 6), and error bars the combination 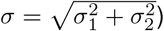 of the standard error of the mean (*σ*_1_) and the mean experimental error (*σ*_2_). Coloured lines: fitted elliptic model. Dark and light grey dotted lines: fitted circular and rotated ellipse model. The inductive shift is approximately sinusoidal, though systematic departures are observed in the data, the shape of which varies with the surround, and is captured by the two elliptic models.

**Figure 7.**
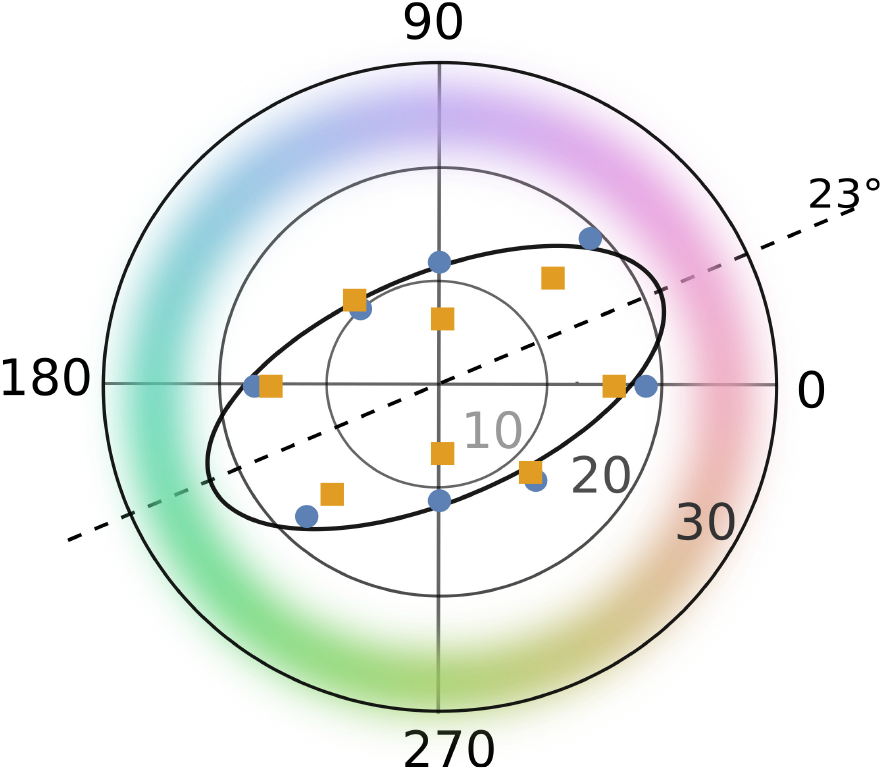
Polar plot reporting the maximal inductive shift Δ*θ* in absolute value, obtained as the average of the two peaks, expressed in degrees, as a function of the angular chromaticity of the surround *θ*_*c*_. Blue dots: Maximal Δ*θ* of each collection of black data points (experimental data) of Fig. 6. Orange squares: Theoretical prediction from Sect. 3.4.1, obtained as the average maximal value of each coloured curve in Fig. 6. Dashed line: Theoretical prediction of the surround producing maximal induction from Sect. 3.4.2. In all three estimations, maximal induction is obtained for surrounds along the purple-green axis.

In Fig. 6, the different surrounds produce induction curves that have slightly different shapes. These differences arise from the combined push produced by the grey surround on the adjustable stimulus, and by the chromatic surround on the target stimulus. If the locus were a circumference in the perceptual coordinates, then the push produced by the grey surround would always be perpendicular to the locus, so it would not affect the shape of the curve Δ*θ*(*θ*_*r*_ − *θ*_*c*_). However, when the locus is an ellipse, the grey surround also contributes with a push, and thereby distorts the curve in a way that depends on the surround chromaticity *θ*_*c*_.

#### 3.4.2 Elliptic 𝒞 with arbitrary orientation

The discrimination ellipses reported by Krauskopf and Gegenfurtner [16] were aligned with the MacLeod-Boynton axes. This implies that those two axes are perceptually perpendicular to one another. However, this may not be the case in other studies, since often, at the very beginning, calibration experiments are performed to determine the plane that the subject under study perceives as iso-luminant. It may well be the case that this plane do not coincide with the plane *L* + *M* = constant. If so, given that experiments typically explore the iso-luminant plane, a variation in *S* and/or *L* − *M* also implies a variation in *L* + *M*. As a consequence, the coordinates *L* − *M* and *S* reported in the study may not coincide with the axes of Macleod and Boynton, and may not be perceptually orthogonal (see Sect. 2.4 for the procedure employed by Klauke and Wachtler [15]). In such case, the perceptual coordinates are related to the experimental ones by the transformation 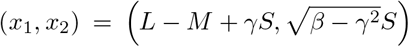, where the parameter *γ* is related to the difference between the inclination of the iso-luminant plane of the tested subject and that of the plane *L* + *M* = constant. A chromatic locus that is circular in the original coordinates is thereby mapped onto an ellipse that is rotated with respect to the vertical axis in an angle φ_0_ given by 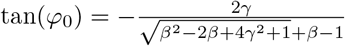.

By fitting the value of *γ*, the match between the theory and the experimental data is improved (grey dotted line in Figs. 6), with *χ*^2^-values of the order of unity (*χ*^2^ = 0.96 per degree of freedom, Table 2), implying a 30% reduction with respect to the un-rotated case. Moreover, the direction of maximal inductive shift is analytically derived to be *θ*_*max*_ = 23°, in excellent agreement with the data (Fig. 7).

Allowing the mapping between the original and the perceptual coordinates to include two fitting parameters *β* and *γ* cannot worsen the fit to the experimental data. To assess the significance of the improvement, a cross-validation procedure was performed, whereby the value of the parameters was fitted with the eight values of *θ*_*c*_ randomly assigned to the eight measurements of experimental data. If the improvement in fit brought about by *β* and *γ* were simply due to the addition of parameters, then the *χ*^2^-value should not differ significantly between the fits obtained with the real and the randomised assignment of *θ*_*c*_. However, the random reordering yields *χ*^2^ = 1.8, which is twice the real value. We therefore conclude that the improvement introduced by the elliptical case with respect to the circular case are consistent with the specific geometry of the experiment.

#### 3.4.3 Fourth-order locus 𝒞

So far, we have considered discrimination ellipses that, in the original coordinates, differed from circumferences in no more than a constant compression factor along a given axis. As a consequence, the mapping between the experimental and the perceptual coordinates was linear, and in the perceptual coordinates, the locus **𝒞** was elliptic. If, however, the discrimination ellipses vary throughout colour space in either size or orientation (or both), the coordinate transformation may be arbitrary, as well as the shape of the locus.

Increasing the number of degrees of freedom in the unknown geometry of the locus is expected to improve the performance of the model in reproducing the data, and thereby, to diminish the falsifiability of the model. Therefore, to avoid overfitting, we only consider geometries which are supported by previous experimental results. Discrimination experiments show increments in perceptual thresholds in the direction *S* of DKL space which are compatible with a quadratic correction in the dependency of perceptual coordinates with the coordinate *S* (Krauskopf and Gegenfurtner [16], Vattuone et al. [38]). When including this correction, the chromatic locus takes the shape of a quartic curve in perceptual coordinates, such as the one shown in Fig. 3, panel B. The explicit parametrisation of the locus is

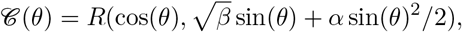

where *α* accounts for the quadratic correction along direction *S*. In the experiment of Klauke and Wachtler [15], the fitted value of *α* is small (*α* = 0.12 ± 0.05) and the improvement in the reduced chi-squared with respect to the case *α* = 0 is only 3% (Table 2), from where the variation of JNDs along direction *S* can be concluded not to be a relevant factor for explaining this data set.

#### 3.4.4 Mean induction for arbitrary geometry

In this section, we extend the analysis to loci of arbitrary shape, as long as they remain simple and closed, and their eccentricity be small. In other words, the departure from the circle that best approximates the locus is assumed to be small (Fig. 8). We calculate the mean inductive shift Δ*θ*(*θ*_*r*_, *θ*_*c*_), when averaged across a collection of surrounds *θ*_*c*_, and also a collection of reference colours *θ*_*r*_, in such a way as to keep the difference *φ* = *θ*_*r*_ − *θ*_*c*_ constant. This operation is equivalent to piling the eight curves of Fig. 6 one on top of another, and averaging them together, as done in Fig. 5. We here prove that the result of this operation coincides with the inductive shift predicted for a circular locus. In other words, to a first-order approximation, the departures from the circular result produced by different surrounds cancel out.

**Figure 8.**
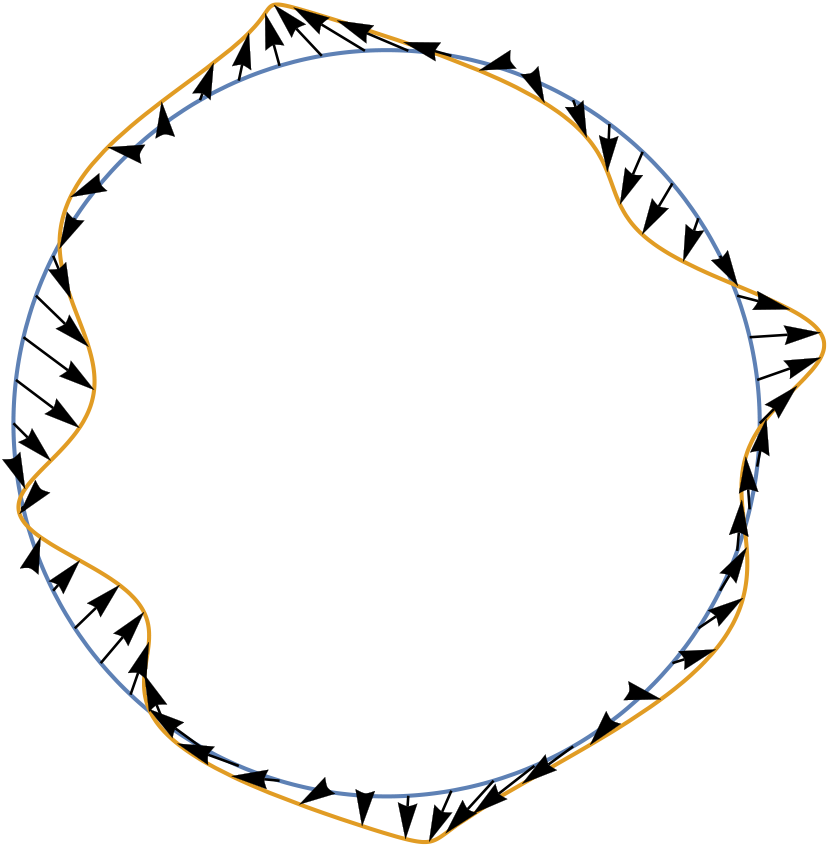
An arbitrary periodic, simple curve **𝒞** (yellow) and the circumference that best approximates it in quadratic norm (blue).

Asymmetric matching experiments are typically designed in such a way that, in the original coordinates, both the set of available colours to produce the match, and the set of explored surrounds, are located on two concentric circumferences. When representing these circumferences in the perceptual coordinates, they no longer appear to be circular. The matching stimuli conform the locus **𝒞**, and it is often the case that the explored surrounds lie on a curve that is proportional to **𝒞**. That is, every surround ***c*** can be written as *c* = *λC*(*θ*_*c*_), for some scalar *λ*.

We start from Eq. 11. The goal is to calculate the average of Δ*θ*(*θ*_*s*_, *θ*_*c*_) across many surrounds *θ*_*c*_, and for each surround, shifting *θ*_*s*_ so that the relative angle *φ* = *θ*_*s*_ − *θ*_*c*_ remain fixed. To a first-order approximation, the result of the average turns out to be equal to the inductive shift of the circular case Δ *θ*°(*φ*), given by the equation 12. Mathematically, this means that

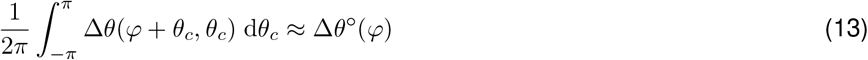

The full argument is developed in the appendix. Here we delineate the main ideas of the proof. First, Eq. 11 is interpreted as a functional equation, which assigns a numeric value for every choice of the curve **𝒞** (*θ*_*s*_). Second, the functional thus defined is shown to have an extreme when evaluated on circumferences. This implies that departures from the circumference have a quadratic effect in the value of the mean chromatic induction. Finally, we explicitly find the circumference that best approximates a given curve **𝒞** (*θ*_*s*_), and show that the effect of the geometry of **𝒞** (*θ*_*s*_) is only quadratic in the mean induction. We therefore conclude that the good agreement shown in 5 between the mean chromatic induction and the prediction with no free parameters made by our model is *independent* of the details of the actual geometry of colour space, as long as the chromatic locus is not too eccentric in that geometry.

### 3.5 Complete model

So far, the comparisons of the model with the experimental data was done with Eq. 11, which had been derived under the approximation that 𝓁 ≫ *κ*, in other words, assuming that the separation between the reference colour *r* and its background *c* was large enough for the inductive shift to have reached its asymptotic value. If this approximation does not hold, then the inductive shift has to be calculated with the full Eq. 8. For that to be possible, the functional shape of **Φ**_***b***_(*r*) is needed. In Vattuone et al. [38], we showed that the inductive function was well described by a monotonically increasing function saturating at |*b* − *r*| ≲ 𝓁, as for example,

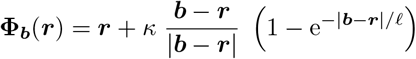

The use of this formula requires to also fit the saturating length 𝓁. Table 2 shows that the fit of the experimental data does not improve by adding this extra parameter, so we conclude that the data collected so far is compatible with the saturation hypothesis.

## 4 Discussion

A long tradition in mathematical neuropsychology has aimed at furnishing sets of concepts with a metric structure, as done for example by von Helmholtz [40], Shepard [28], Treves [35], Constantinescu et al. [4], among others. The utility of this endeavour has recently been questioned as a rather futile intellectual exercise [3, 10]. Contrasting with this somewhat skeptic perspective, here we have provided evidence that at least for some conceptual spaces — specifically, the space of colours — insight is indeed gained by endowing colours with a metric structure. Our analysis was based on the following assumptions: (1) The space of colours can be described by a Riemannian metric. (2) A set of coordinates exists in which the metric is nearly Euclidean. (3) Chromatic induction is isotropic and homogeneous in these coordinates. (4) Chromatic induction saturates for long distances. These assumptions allowed us to derive, with no free parameters, the chromatic induction averaged along a closed locus of arbitrary shape. The prediction matches the existing experimental data even though at first glance there were no evident symmetries in the data, because the experiments had been designed in a system of coordinates in which chromatic induction was neither rotationally nor translationally invariant.

The previous studies suggested that the inhomogeneities and anisotropies in colour space observed in asymmetric matching tasks could be the result of natural selection favouring a particularly large inductive shift for blue stimuli surrounded by purple, or yellow stimuli surrounded by green. The present study shows that these asymmetries disappear with an adequate choice of the metric of colour space. Yet, we do not deny the possibility that evolution may have selected observers with large induction in specific regions of colour space. Our observation only points out that the effect can be entirely absorbed by the metric, because ultimately, the assessment of whether a shift is large or small is unavoidably linked to the choice of a notion of distance. Therefore, if there is a selection pressure for magnifying the representation of certain regions of colour space, the pressure probably acts at the level of the metric, and not on chromatic induction specifically. The metric describes the internal representation of colour, and as such, it influences many more phenomena than just induction: it also affects detection and discrimination thresholds, colour naming, and probably other phenomena too. Therefore, it would be valuable if future work were directed to assess whether the asymmetries observed in these other tasks can also be derived from the perceptual metric. Some previous experiments have already addressed the issue of detection and discrimination thresholds [38].

When the metric is flexible enough to describe JNDs that are elliptic in the experimental coordinates, or equivalently, loci that are elliptic in the perceptual coordinates, the theory predicts the inductive shift quite successfully (Sects. 3.4.1 and 3.4.2). At most two parameters are fitted for all the surrounds, and many of the peculiar bends in the curve Δ*θ*(*θ*_*r*_ − *θ*_*c*_) are explained by the non-aligned pushes produced by the grey and the chromatic surrounds. Increasing the number of parameters describing the metric does not improve the fit.

The success of the model raises the question of whether the geometric structure of colour space is implemented at the physiological level as some function of the activity of populations of neurons representing colour, or whether it is merely a mathematical gimmick that provides an effective description that has no physiological correlate. To assess whether the first hypothesis holds, as future work we plan to evaluate a physiologically plausible model of the transformations involved in the representation of colour in the early stages of visual processing, including a description of induction in which horizontal connections instantiate the perceptual shift produced by surrounds. The model, while realistic, should yield a representation of colour that is invariant to rotations and displacements in the perceptual coordinates. If, alternatively, the Riemannian metric is only an effective description that has no correlate at the physiological level, we should find a theoretical framework that explains how a symmetric model can predict experimental results produced by a system that contains no inherent symmetries in its mechanistic realisation.

## Acknowledgements

This work was supported by Consejo Nacional de Investigaciones Científicas y Técnicas, Comisión Nacional de Energía Atómica and Agencia Nacional de Promoción Científica y Tecnológica of Argentina. We are thankful to Thomas Wachtler, for his comments and insight.

## 5 Appendix

### 5.1 Circular aproximation

In this section we prove Eq. 13. We fist define the functional

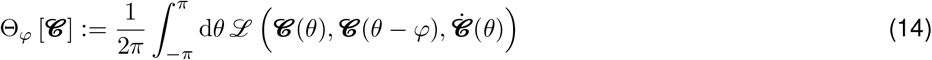

where the Lagrangian ℒ is

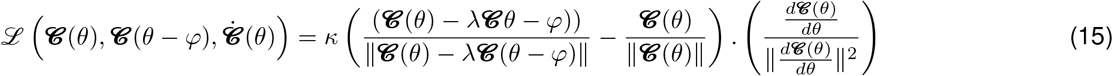

is the inductive shift Δ*θ* of Eq. 11 but seen as a function of the chromatic curve. Then Θ_*φ*_ [**𝒞**] is the mean inductive shift when averaged over the chromaticity of the surround. To prove equation 13 we

1. find the circumference **𝒞** ° that best approximates the curve **𝒞** (*θ*),
2. expand Θ_*φ*_ [**𝒞**] around **𝒞** °, and
3. prove that the linear term in the expansion vanishes.

To achieve the first step, we fix the origin of coordinates at the barycentre of **𝒞** (*θ*), which implies that the integral 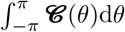 vanishes. Next, we write

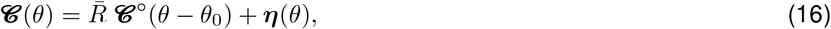

where **𝒞** °(*θ*) = (cos(*θ*), sin(*θ*)) is the unit circumference and 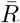 and *θ*_0_ are unkown parameters. We want to find the values for 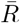 and *θ*_0_ for which the quadratic deviation

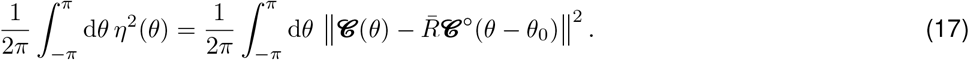

is minimal. Requiring the partial derivates of the quadratic deviation with respect to *R* and *θ*_0_ to vanish,

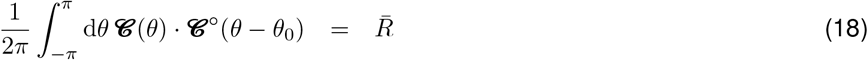

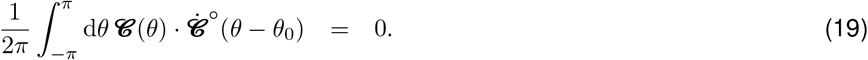

The first expression yields 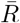 explicitly. The second, is an implicit equation for *θ*_0_, and since the value of the left-hand side changes sign when *θ*_0_ increases in *π*, continuity guarantees that there is always a solution to the equation.

Once the values of 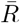 and *θ*_0_ have been found, if Eq. 16 is inserted in Eqs. 18 and 19

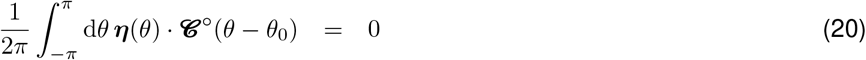

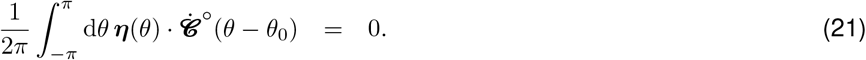

Step 2 can be approached with the calculus of variations, with which Θ_*φ*_ [**𝒞**] can be approximated byD

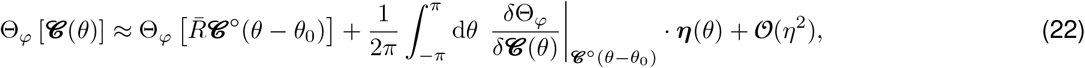

where the functional derivative yields the Euler-Lagrange equations

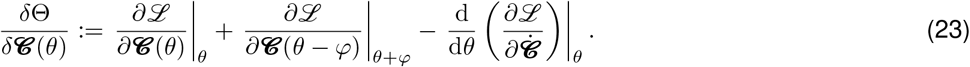

The Lagrangian depends on the curve evaluated in two different values (*θ* and *φ* + *θ*), so the Euler-Lagrange equations contain one term for each value. Since **𝒞** has two components (*C*_1_, *C*_2_), the derivative *∂/∂* **𝒞** is

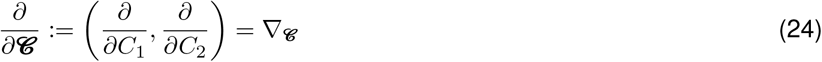

The calculation is long but straightforward, and the final results is

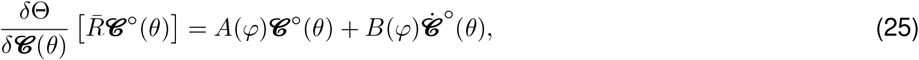

Where

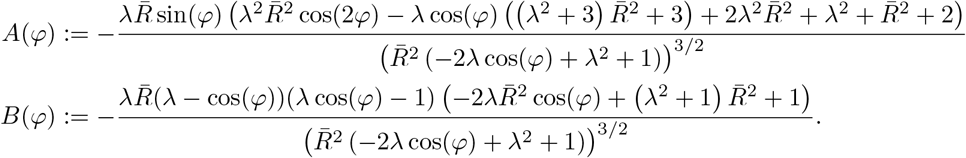

Replacing this result in Eq. 22,

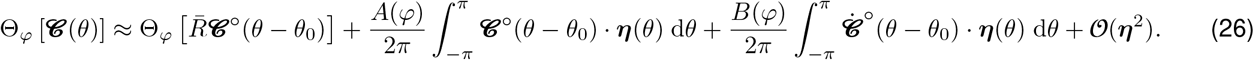

Equations 20 and 21 imply that both integrals in Eq. 26 vanish, proving the validity of step 3, and yielding

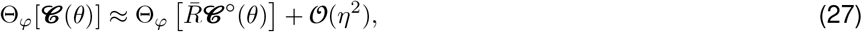

which was the goal of this appendix.

### 5.2 Summary of models, parameters and goodness of fit

